# Reactive Oxygen Species Modulate Activity-Dependent AMPA Receptor Transport in *C. elegans*

**DOI:** 10.1101/2020.04.20.051904

**Authors:** Rachel L. Doser, Gregory C. Amberg, Frederic J. Hoerndli

**Affiliations:** Department of Biomedical Science, Colorado State University, Fort Collins, Colorado 80523

## Abstract

The AMPA subtype of synaptic glutamate receptors (AMPARs) play an essential role in cognition. Their function, numbers and change at synapses during synaptic plasticity, is tightly regulated by neuronal activity. Although we know that long-distance transport of AMPARs is essential for this regulation, we don’t understand the regulatory mechanisms of it. Neuronal transmission is a metabolically demanding process in which ATP consumption and production are tightly coupled and regulated. Aerobic ATP synthesis unavoidably produces reactive oxygen species (ROS), such as hydrogen peroxide, which are known modulators of calcium signaling. Although a role for calcium signaling in AMPAR transport has been described, there is little understanding of the mechanisms involved and no known link to physiological ROS signaling. Here, using real-time *in vivo* imaging of AMPAR transport in the intact *C. elegans* nervous system, we demonstrate that long-distance synaptic AMPAR transport is bidirectionally regulated by calcium influx and activation of calcium/calmodulin-dependent protein kinase II. Quantifying *in vivo* calcium dynamics revealed that modest, physiological increases in ROS decrease calcium transients in *C. elegans* glutamatergic neurons. By combining genetic and pharmacological manipulation of ROS levels and calcium influx, we reveal a mechanism in which physiological increases in ROS cause a decrease in synaptic AMPAR transport and delivery by modulating activity-dependent calcium signaling. Taken together, our results identify a novel role for oxidant signaling in the regulation of synaptic AMPAR transport and delivery, which in turn could be critical for coupling the metabolic demands of neuronal activity with excitatory neurotransmission.

**SIGNIFICANCE AND IMPACT:** Synaptic AMPARs are critical for excitatory synaptic transmission. The disruption of their synaptic localization and numbers is associated with numerous psychiatric, neurological, and neurodegenerative conditions. However, very little is known about the regulatory mechanisms controlling transport and delivery of AMPAR to synapses. Here, we describe a novel physiological signaling mechanism in which ROS, such as hydrogen peroxide, modulate AMPAR transport by modifying activity-dependent calcium influx. Our findings provide the first evidence in support of a mechanistic link between physiological ROS signaling, AMPAR transport, localization, and excitatory transmission. Of potential therapeutic importance, dysregulation of intracellular calcium and ROS signaling is implicated in the pathogenesis of several neurodegenerative disorders including Alzheimer’s and Parkinson’s disease.

## INTRODUCTION

The alpha-amino-3-hydroxy-5-methyl-4-isoxazole-propionic acid (AMPA) subtype of glutamate receptors (AMPARs) are essential for fast excitatory synaptic transmission (Ashby et al., 2008).The number of AMPARs at the synaptic surface is a key determinant of synaptic efficacy and is the result of a dynamic equilibrium between the number of receptors in intracellular pools and at the synaptic surface (Rosendale et al., 2003; Groc et al., 2009; Henley and Wilkinson, 2013). Although a few AMPARs can be synthesized locally (Hanus et al., 2016), the vast majority of AMPARs are synthesized in the neuronal soma, often far away from synapses, and are trafficked in a complex, multistep process to dendrites and synapses (Hanus et al., 2016; Henley and Wilkinson, 2016; Brechet et al., 2017). This includes intracellular transport by molecular motors (Kim and Lisman, 2001; Setou et al., 2002; Hoerndli et al., 2013; Esteves da Silva et al., 2015; Hangen et al., 2018) exo- and endocytosis (Ehlers, 2000; Yudowski et al., 2007) as well as surface diffusion dynamics (Choquet and Triller, 2013). Intracellular AMPAR transport between different cellular pools of AMPARs is the least understood of these steps. Recent studies have revealed that long-distance transport is highly regulated by neuronal activity (Esteves da Silva et al., 2015; Hoerndli et al., 2015; Hangen et al., 2018). However, the regulatory mechanisms of activity-dependent AMPAR transport are still poorly defined.

Neuronal activity is associated with increased energy demands that is largely fulfilled by mitochondrial ATP production (Hall et al., 2012), which concurrently produces reactive oxygen species (ROS; Halliwell, 1992). The main ROS species are hydrogen peroxide (H_2_O_2_), the superoxide anion (O_2_^-^) and the hydroxyl radical (HO^-^) (Halliwell, 1992). Previous studies have shown that ROS can affect calcium signaling mediated by N-methyl-D-aspartate (NMDA) glutamate receptors, voltage-gated calcium channels (VGCCs) and calcium release from the endoplasmic reticulum (ER) (Akaishi et al., 2004; Amberg et al., 2010; Todorovic and Jevtovic-Todorovic, 2014; Hidalgo and Arias-Cavieres, 2016). Interestingly, the effect of ROS varies widely depending on dosage, cell type and model system used (Hidalgo and Arias-Cavieres, 2016; Wilson et al., 2018). This is illustrated by the fact that long-term potentiation (LTP) is disrupted by elevated ROS (Bliss and Collingridge, 1993; Klann, 1998; Kamsler and Segal, 2003) as well as depletion of ROS (Kishida and Klann, 2006). Thus, the literature supports a link between ROS signaling and changes in neuronal excitability. However, whether this is due to ROS modulation of calcium signaling remains uncertain. In particular, there is a lack of direct evidence for the roles of physiological ROS on *in vivo* neuronal calcium signaling. The transparent model *C. elegans* is well-suited to study the effects of ROS on calcium signaling in neurons *in vivo* where circuits remain intact. The unique combination of genetic manipulations, calcium sensors and pharmacological agents allow us to investigate the targets and effects of physiological ROS signaling *in vivo*. Finally, because of the role of activity-dependent calcium signaling in AMPAR transport, we propose a novel link between ROS signaling and the regulation of AMPAR transport.

In this study, we start to address this question using the transparent model *C. elegans*. Single neuron expression of SEP::mCherry::GLR-1 (the *C. elegans* homologue of the AMPAR subunit GluA1) and the calcium sensor GCAMP6f enabled us to quantify and characterize GLR-1 transport as well as calcium influx *in vivo*, in real time. Together with genetic and pharmacological manipulation of VGCC activity and ROS levels, our results show that AMPAR transport is directly regulated by activity-dependent calcium influx. We also find that physiological increases in ROS levels decreases calcium influx and, as a result, AMPAR transport, delivery and exocytosis. We further show that the targets of increased ROS are specific and involve L-type VGCC-dependent calcium signaling upstream of calcium/calmodulin-dependent protein kinase II (CaMKII) activation. Altogether, our results suggest a mechanism by which physiological ROS signaling acts as a negative feedback mechanism regulating excitatory glutamatergic transmission by decreasing activity-dependent calcium influx and subsequent AMPAR transport.

## MATERIALS AND METHODS

### Strains

*C. elegans* strains were maintained on nematode growth media (NGM) and fed with the *E. coli* strain OP50 at 20°C (Brenner, 2003). Strains used in these experiments contained the following alleles:

**Table 1:** List of genetic alleles the corresponding gene, the effect of the mutation and reported functional change along with original references mentioning the allele.

| Gene | Allele | Mutation | Functional Change | References |
| --- | --- | --- | --- | --- |
| *glr-1* | *ky176* | Premature stop | Truncated, unfunctional receptor | (Maricq et al., 1995) |
| *egl-19* | *n582* (rf) | Missense mutation in S4 domain | No calcium spikes | (Trent et al., 1983; Liu et al., 2018) |
|  | *n2368* (gf) | Missense mutation in IS6 | Delayed inactivation, prolonged calcium spikes | (Lee, 1997; Laine et al., 2014) |
| *unc-43* | *n498n1186* (lf) | Premature stop | Protein null | (Park and Horvitz, 1986; Reiner et al., 1999; Umemura et al., 2005) |
|  | *n498sd* (gf) | Missense mutation in the active site | Partial calcium-independent, constitutive activation |  |
| *ctl-2* | *ok1137* (lf) | ~1kb deletion | Protein null | (Spiró et al., 2012) |
| *lin-15* | *n765ts* | Frameshift mutation | Protein null | (Kim and Horvitz, 1990) |

### Cloning

The catalase gene (*ctl-2*) was cloned from *C. elegans* cDNA using the forward primer 5’-GGGGACAAGTTTGTACAAAAAAGCAGGCTATGCCAAACGATCCATCGGA-3’ and reverse primer 5’-GGGGACCACTTTGTACAAGAAAGCTGGGTCGATATGAGAGCGA GCCTGTTTC-3’ designed on ApE (v.2.0.60). Using the gateway recombination cloning technique (Invitrogen), the *ctl-2* gene was positioned behind *flp-18*, an AVA specific promoter (Schmitt et al., 2012), and followed by an eGFP-containing 5’ UTR (pGH112, Erik Jorgensen) in the destination vector pCFJ150 (Erik Jorgensen, Addgene plasmid #19329). The GCaMP6f-containing plasmid (*Prig-3*::GCaMP6f::*unc-54*) was a gift from Attila Stetak.

### Transgenic strains

For visualization of GLR-1, we used akIs201, an integrated array containing *Prig-3*::SEP::mCherry::*glr-1* (Hoerndli et al., 2015), which allowed for cell-specific expression of dual-tagged GLR-1 in the AVA glutamatergic interneurons. Transgenic strains were created by microinjection of *lin-15(n765ts)* worms with plasmids containing *lin-15(+)* to allow for phenotypic rescue of transgenic strains. The above plasmids were used to create strains expressing the following extrachromosomal arrays: csfEx65 [*Pflp-18*::*ctl-2*::eGFP + *Prig3*::*glr-1*::mCherry] and csfEx62 [*Prig-3*::GCaMP6f::*unc-54*].

### Confocal microscopy

Imaging was carried out on a spinning disc confocal microscope (Olympus IX83) equipped with 488 nm and 561 nm excitation lasers (Andor ILE Laser Combiner). Images were captured using an Andor iXon Ultra EMCCD camera through either a 10x/0.40 or a 100x/1.40 oil objective (Olympus). Devices were controlled remotely for image acquisition using MetaMorph 7.10.1 (Molecular Devices).

### Transport imaging and analysis

All transport imaging was conducted on strains containing akIs201 in the *glr-1* null background (*ky176*). One-day-old adults from these strains were mounted on a 10% agarose pad with 1.6 µL of a mixture containing equal measures of polystyrene beads (Polybead, Polysciences Inc.) and 30 mM muscimol (MP Biomedicals). The worm was positioned to allow for the AVA interneuron to be in close proximity to the coverslip through which the AVA neurite would be imaged. Once the AVA was located using the 100x objective and a 561 nm excitation laser, a proximal section of the AVA was photobleached using a 3 W, 488 nm Coherent solid-state laser (Genesis MX MTM) set to 0.5 W output and a 1 s pulse time. The photobleaching laser was targeted to a defined portion of AVA using a Mosaic II digital mirror device (Andor Mosaic 3) controlled through MetaMorph. Immediately following photobleaching, a 500-frame image stream was collected in a single z-plane with the 561 nm excitation laser with a 100 ms exposure time. Kymographs were generated using the Kymograph tool in MetaMorph with a 20 pixel line width as previously reported (Hoerndli et al., 2013). Transport quantification, stops and velocities were analyzed using the ImageJ plugin KymoAnalyzer (Neumann et al., 2017).

### FRAP

Strains containing *akIs201* were mounted for imaging as described above. After locating the AVA, an imaging region was determined, and its location was recorded using MetaMorph’s stage position memory function. Then, an image stack was acquired using the 561 nm then the 488 nm excitation laser (500 ms exposure) around the AVA process (20 images were taken every 0.25 µm starting 2.5 µm below to 2.5 µm above the process). A ∼80 µm proximal and distal section of the AVA flanking the imaging region was photobleached using the same photobleaching settings as previously described. The imaging region was then photobleached. Immediately after, two image stacks (with 561 nm then 488 nm excitation) was acquired for the 0-minute time point. This was repeated at 2, 4, 8 and 16 minutes following the photobleaching of the imaging region. Image stacks from all time points were converted to maximum projections using MetaMorph’s stack arithmetic function. The average fluorescence in the imaging region at each time point was analyzed using the region measurement tool in ImageJ 1.51s (Java 1.8). The background fluorescence (i.e. outside of the AVA) from each maximum projection was then subtracted from the average fluorescence of the imaging region. The resulting fluorescence from the maximum projections 0 minutes following photobleaching was subtracted from the fluorescence values of all subsequent time points. These values were then compared to those from images taken before photobleaching to determine the percent of signal recovery for each channel within the imaging region.

### *In vivo* calcium imaging

All strains used for calcium imaging experiments contained the extrachromosomal array csfEx62 expressing GCAMP6f in the AVA interneurons in the *lin-15(n765ts)* genetic background. Eight to ten one-day-old adult animals with the array were selected and placed on a 10% agar pad with 2 μL of M9 buffer. *C. elegans* animals were thus constrained but not immobilized similar to when placed in a microfluidics chamber (Chronis et al., 2007). Animals were imaged using the 10x objective on the spinning disc confocal. A 60-second image stream consisting of 240 images with a 250 ms exposure time was acquired using the 488 nm excitation laser. During imaging, animals spontaneously attempt reversals, which is correlated with activation of the AVA, calcium influx (Ben Arous et al., 2010) and thus changes in GCaMP6f fluorescence in our strains. We report the total activity or total influx of calcium during each 60-second stream (Figure 2 and extended Figure 2-1). This was calculated using the following approach.

For each frame, the maximum fluorescence (F_(t)_) was quantified using MetaMorph’s region measurement tool by manually defining the region of the image stream containing the AVA cell bodies. Attempts at reversals exhibited large calcium transients whereas attempts at forward movement or absence of movement was correlated with only small variations considered as basal fluctuations. The baseline (F_min_) was defined as the average GCAMP6f signal when worms are immobile or during forward movement as this was previously shown not to activate AVA (Ben Arous et al., 2010). GCAMP6f signal during this time was observed to be within 30% of the overall minimum value of GCAMP6f. Maximum fluorescence values for each frame was imported into a customized Excel document containing modules created with Excel’s visual basic editor. One module calculates the average baseline (F_min_) by averaging all values within 30% of lowest value. This value was used to determine the ΔF (F_(t)_ -F_min_) for each frame normalized to the average baseline (ΔF/F_min_; Larsch et al., 2013). Total activity was defined as the sum of all F_(t)_ values greater than F_min_ (Green area below the curve, Extended Figure 2_1) normalized to the average baseline (average values below dotted line, Extended Figure 2_1).

### Spontaneous reversal quantification

The spontaneous reversal rate (reversal/min) was quantified using the semi-automated tracking system of the WormLab System (MBF Bioscience). Briefly, for each trial, five to eight one-day-old adult worms were selected off plates with bacterial food and transferred on food free plates twice with a rest of ∼2 minutes after each transfer. Spontaneous locomotion was then recorded for 1 minute on the second plate and the video analyzed blinded to genotype. After detection of all animals and tracks by the semi-automatic tracking system, all tracks were manually verified and corrected if necessary, still blind to genotype. N2, wild-type animal locomotion and reversal rates were consistent with previously reported values: 3.78 ± 0.26 reversal/minute compared to reported ∼ 4 ± 0.2 (Monteiro et al., 2012).

### Hydrogen peroxide and nemadipine treatments

Worms were acutely treated with H_2_O_2_ by placing the worm in 1.6 uL of solution containing the appropriate concentration of H_2_O_2_ with either 15 mM Muscimol and polystyrene beads (for transport imaging) or M9 buffer (for calcium imaging). Approximately 5 minutes following the beginning of the treatment, image streams were acquired. Physiological intracellular concentrations of H_2_O_2_ range from 10-100 nM in vertebrate neurons (Sies, 2017), so 10, 50 and 100 nM were used for our experiments.

A 0.5 mM stock of nemadipine (Enzo Life Sciences) was dissolved in 0.1% DMSO and the concentration was adjusted to 10 μM (a concentration reported to decrease calcium influx in exposed neurons *in vivo* by ∼70%; Larsch et al., 2013) with M9 buffer and Op50 liquid culture immediately before a 30-minute treatment. During the treatments, one-day-old adult wild-type worms were placed into 1.5 mL Eppendorf tubes containing either control media or the pharmacological agent and placed on a rocker for oxygenation. The worms were then pipetted onto fresh NGM/Op50 plates immediately before being moved to an agar pad for imaging.

### Image presentation and data analysis

All images were acquired under non-saturating conditions. Quantification of GLR-1 transport, FRAP and calcium imaging is described in the appropriate sections above. Representative images shown were chosen and processed following analysis only to the extent necessary to appreciate the corresponding quantifications. Image processing (RGB colors, cropping, adjustment of brightness and contrast) was performed in Photoshop (21.1.1). For FRAP images, the mCherry signal of SEP::mCherry::GLR-1 is shown in magenta for colorblind vision. This was achieved in Photoshop by duplicating the 561 nm information to create the red and blue channels of the RGB image which was then merged to create the magenta image as published previously (Hoerndli et al., 2013). All images in each panel were identically processed.

### Experimental Design and statistical analysis

All experiments were performed using one day old adult hermaphrodite *C. elegans* animals as determined by a single row of eggs and picking as precisely identifiable L4 stage larva 24 hours before imaging and behavior experiments. All mutant strains were back-crossed with N2 wild-type animals at least 2x. All imaging reagents such as SEP::mCherry::GLR-1 and GCAMP6f were crossed into strains carrying genetic mutations in the exact same way, verifying the presence of the knock-out *glr-1* allele *ky176* and genetic mutations using PCR genotyping on at least 2 generations. Primer sequences are available on request.

For statistical analysis, all datasets were screened for outliers using a Thompson Tau test. For datasets including only two experimental groups, statistical significance was tested using a two-tailed student’s t-test. For datasets comparing more than two experimental groups, a one-way Brown-Forsythe ANOVA with a Dunnett’s correction for multiple comparisons was used. FRAP differences between groups was determined using an extra sum-of-squares F-test of the nonlinear regression fit to the data. All statistical analyses were performed using Prism 8 software. Statistical details, numbers of ROIs and animals analyzed, and *p* values are indicated in detail in each Results section. Data are presented as mean ± SEM unless otherwise stated.

### Code/Software

Code for custom Excel modules used for analysis of calcium imaging is available upon request.

## RESULTS

*C. elegans* are a useful model for studying long-distance AMPAR transport dynamics *in vivo*. Here we use a dual tagged AMPAR subunit, SEP::mCherry::GLR-1, expressed in a single pair of glutamatergic neurons (AVA) to analyze how transport, delivery and exocytosis of GLR-1 are modulated by calcium influx and reactive oxygen species (ROS). AVA are long, ventrally running unipolar interneurons with cell bodies in the head of the animal. These neurons express AMPA and NMDA subtypes of glutamate receptors (Maricq et al., 1995) and play an essential role in associative olfactory memory in *C. elegans* (Stetak et al., 2009). To reveal dim GLR-1 transport events, we used a photobleaching approach combined with streaming imaging of the mCherry signal to visualize GLR-1 transport in the proximal region of AVA (see Figure 1A). Both anterograde (Figure 1A, blue arrowheads) and retrograde (Figure 1A, fuchsia arrowheads) transport events can be visualized as single particles advancing in opposite directions at different time points (Figure 1A, timepoint images 1-3). The trajectories of these transport events can be visualized and analyzed in a kymograph representing their position on x-axis and time on the y-axis (Figure 1A, bottom right). The total amount of GLR-1 transport as well as velocities and stopping of transport events can be quantified using these kymographs. In wild-type animals, the number of transport events as well as the average anterograde velocity obtained in our hands is similar to what we reported previously and has been reported for vertebrate AMPAR transport in hippocampal neurons (Ju et al., 2004; Hoerndli et al., 2015; Hangen et al., 2018).

**Figure 1:**
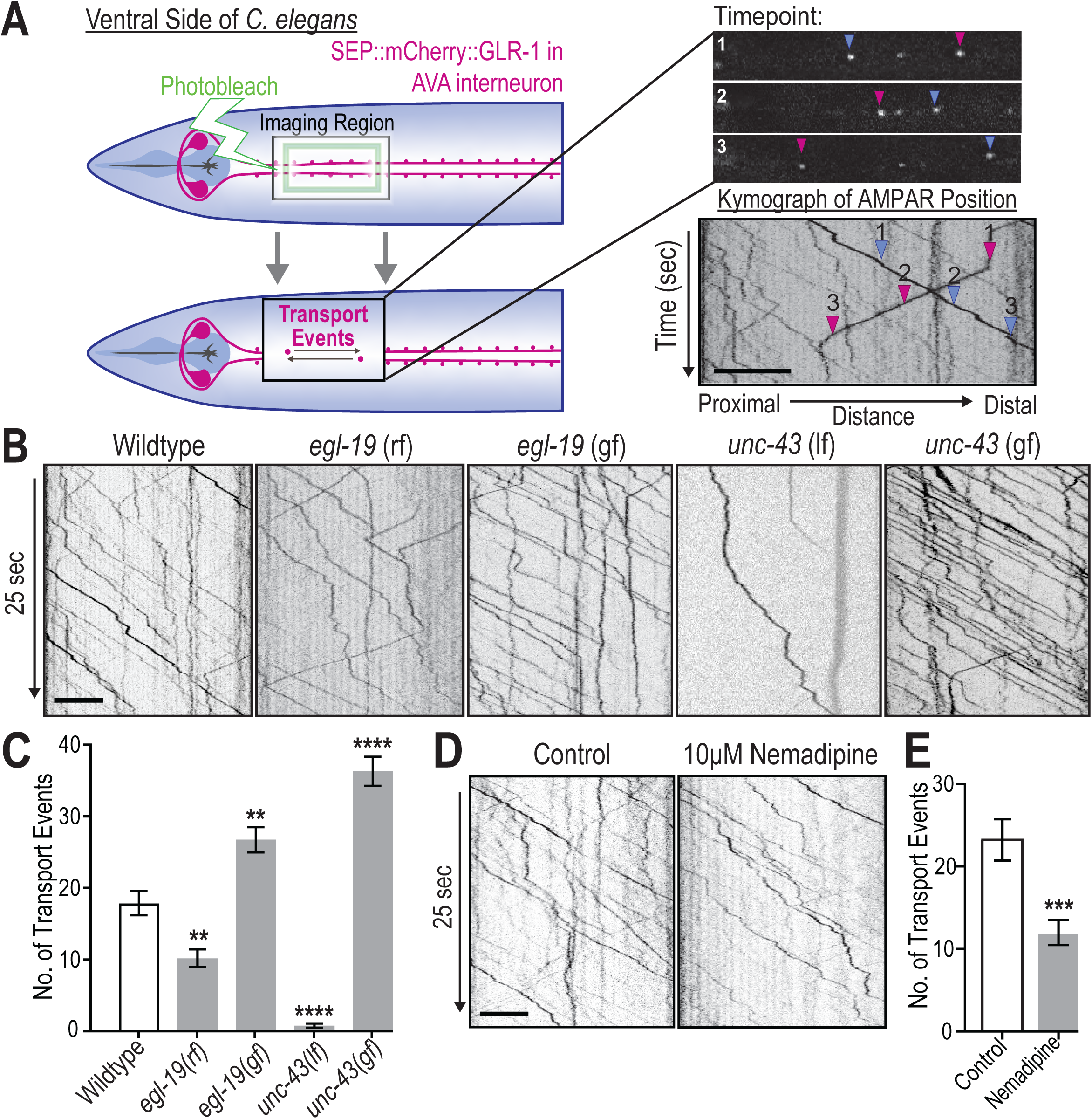
Activity-dependent calcium signaling regulates GLR-1 transport. A) Left, Diagram illustrating the location and procedure for *in vivo* imaging of mCherry::SEP::GLR-1 in AVA. Top right, representative images at three timepoints showing positions of AMPAR-containing vesicles as they progress away from the cell body (blue arrowhead) and toward the cell body (fuchsia arrowhead). Bottom right, a kymograph displaying the position (y-axis) of AMPAR-containing vesicles over time (x-axis). Arrowheads indicate position of vesicles in kymograph that correspond to representative images at timepoints 1-3. Scale bar=5 μm. B) 25 seconds of representative kymographs from wild-type, *egl-19* reduced-function (rf) mutant, *egl-19* gain-of-function (gf) mutant, *unc-43* loss-of-function (lf) mutant, *unc-43* gain-of-function (gf) mutant groups. Scale bar=5 μm. C) Total transport (anterograde and retrograde events) quantified from kymographs representative of a 50 second image stream (n>14 worms for each group; **: p=0.036, ****: p<0.00001 compared to wild-type). D) 25 seconds of representative kymographs of control and 10 μM nemadipine treated wild-type worms. Scale bar=5 μm. E) Total GLR-1 transport as quantified from full-length (50 s) kymographs from control (n=12) and nemadipine treated wild-type worms (n=14, ***: p=0.005 compared to untreated). Error bars for all bar graphs represent standard error of the mean (SEM)

### Activity-dependent calcium signaling regulates AMPAR transport

Recently, we and others have shown that long distance AMPAR transport is regulated dynamically by neuronal activity. Although studies have shown that CaMKII activation is required for activity-dependent AMPAR transport, the exact signaling pathways leading to CaMKII activation are still unclear (Hayashi et al., 2000; Hoerndli et al., 2015; Hangen et al., 2018). In *C. elegans* neurons, the majority of neuronal depolarization is achieved by VGCC, specifically by L-type VGCC (L-VGCC), due to the absence of voltage-gated sodium channels (Serrano-Saiz et al., 2013). *C. elegans* animals expressing VGCCs with reduced calcium conductance have altered synaptic distribution (Rongo C and Kaplan J, 1999) and diminished AMPAR transport (Hoerndli et al., 2015). A necessary next step in understanding the regulation of long-distance transport of AMPARs to and from synapses is to determine if this process is directly regulated by calcium influx leading to CaMKII activation. If this is the case, then we would expect transport characteristics, such as export from the soma as well as transport velocities and pausing, to correlate with activity-dependent changes in cellular calcium levels.

To test this hypothesis, we took a genetic approach using strains with a reduced-or gain-of-function mutation in *egl-19*, the sole L-VGCC gene in *C. elegans*, leading respectively to a decrease or increase in calcium conductance (Liu et al., 2018). In the *egl-19* reduced-function (rf) mutant, GLR-1 transport out of the cell body was significantly decreased (10.2 ± 1.2, mean ± s.e.m., transport events per kymograph, n=16, p=0.0036, Figure 1C) compared to wild-type (17.9 ± 1.7 events, n=19). Conversely, in the *egl-19* gain-of-function (gf) mutant, GLR-1 transport was upregulated (26.7 ± 1.7 events, n=17, p=0.0037, Figure 1B and 1C). To ensure that these changes in transport are in fact due to altered calcium influx, we acutely treated with the L-type-specific VGCC blocker, nemadipine (Kwok et al., 2006). A 30 minute pre-treatment with 10 μM nemadipine caused a decrease in total GLR-1 transport (12.0 ± 1.5 events, n=14, p=0.0005), in comparison to untreated controls (23.4 ± 2.5 events, n=12, Figure 1D and E) comparable to *egl-19(rf)* (10.2 ± 1.2, Figure 1C). Both nemadipine treatment and *egl-19(rf)* showed a similar reduction in total calcium influx in AVA as measured by GCAMP6f (data not shown). These results indicate that not only is calcium influx through VGCCs required (Hoerndli et al., 2015), but also that neuronal calcium levels directly and bidirectionally regulate AMPAR export to and from the cell body.

To better understand how AMPAR transport trajectories are impacted by activity-dependent calcium influx, we quantified the velocity and stop frequency of individual GLR-1-containing vesicles. We found that the increased calcium influx in *egl-19(gf)* mutants results in a faster anterograde instantaneous velocity (1.37 ± 0.03 μm/sec, n=87 events, p=0.003) compared to wild-type animals (1.23 ± 0.02 μm/sec, n=90, Figure 1_1A). The distribution frequency of these velocities in each group revealed that the increase in *egl-19(gf)* is due to more vesicles traveling at higher speeds (1.6-2.0 μm/sec, Figure 1_1C). Retrograde transport velocities, however, were not significantly changed in the *egl-19(gf)* mutants (Figure 1_1A and 1_1D). In addition, we observed a significant increase in the percent time spent paused for vesicles moving in either direction in *egl-19(gf)* (29.9% ± 2.7) compared to wild-type (20.9% ± 1.8, p=0.043, n>95, Figure 1_1B). Decreased calcium influx in *egl-19*(rf) mutants decreases instantaneous velocity of anterograde GLR-1 transport (1.08 ± 1.1 μm/sec, n=50, p=0.0037), but surprisingly had the opposite effect on instantaneous velocity of retrograde transport (1.4 ± 0.06 μm/sec, n=59, compared to 0.9 ± 0.06 μm/sec in wild-type, n=44, p<0.0001, Figure 1_1A). The percent time vesicles moving in either direction spent paused was also drastically decreased in *egl-19(lf)* mutants (paused 10.5% ± 1.5 of the time, p=0.0011, n=79, Extended Figure 1_1B). These data demonstrate that calcium influx directly regulates the amount of GLR-1 transport as well as differentially regulates the dynamics of anterograde and retrograde transport events.

Since CaMKII activity and/or AMPAR phosphorylation by CaMKII is required for normal AMPAR transport (Hoerndli et al., 2015; Hangen et al., 2018), we sought to determine if and to what degree CaMKII activation regulates AMPAR transport. For these experiments, we used strains harboring genetic loss- and gain-of-function mutations of UNC-43, the sole *C. elegans* ortholog of CaMKII. The *unc-43* loss-of-function mutation (*unc-43(lf))* leads to a complete loss of UNC-43 (Reiner et al., 1999), whereas the gain-of-function (*unc-43(gf)*) allele causes partial calcium-independent, constitutive activation of UNC-43 (Umemura, 2005). Animals with *unc-43(lf)* showed a drastic decrease in GLR-1 transport (0.78 ± 0.30, n=14, p<0.0001) whereas animals with *unc-43(gf)* showed a dramatic increase in GLR-1 transport (36.3 ± 2.04 events, n=14, p<0.0001, Figure 1B and C) compared to wild-type. Interestingly, in *unc-43(gf)* mutants, instantaneous transport velocity was unchanged but the percentage of time all transport vesicles spent stopped was drastically decreased to nearly 0% (vesicles stopped 0.06% ± 0.49 of the time, n=58, compared to 20.9% ± 1.8 in wild-type, n=95, p<0.0001, Extended Figure 1_1A and B). We were unable to quantify velocities and stop frequency of GLR-1 transport in *unc-43(lf)* due to the low numbers of transport events per kymograph (less than one per kymograph on average).

Collectively, these results suggest that the amount of calcium influx through L-VGCCs along the neuronal process impacts anterograde transport velocity. Additionally, this calcium influx partially regulates the frequency and duration in which GLR-1 containing vesicles stop along the neuronal process. The drastic decrease in stop frequency in *unc-43*(gf) mutants whose CaMKII activity is independent of calcium influx suggests that stop frequency of GLR-1 transport could be mediated by calcium-dependent changes in CaMKII activity. Taken together, these results provide evidence for a model in which calcium influx through L-VGCCs and CaMKII activation regulate AMPAR transport into and out of the cell body as well as in the dendritic process in an activity-dependent manner. These findings advance our understanding of AMPAR transport by delineating a mechanism in which neuronal activity up- and downregulates the quantity and dynamics of AMPAR transport.

### Increasing ROS levels within physiological range modulates activity-dependent calcium influx

A growing field of evidence shows that calcium-dependent signaling is modified or co-regulated by ROS (Görlach et al., 2015). Additionally, several studies have shown that neural excitability and synaptic plasticity are modified by ROS signaling (Yermolaieva et al., 2000; Kishida and Klann, 2006). More recently, a few studies have shown that function of VGCCs, including the L-type, are altered by increases in ROS above physiological concentrations (Todorovic and Jevtovic-Todorovic, 2014; Dang et al., 2018). However, we have a poor understanding of if and how physiological ROS signaling impacts activity-dependent fluctuations in neuronal calcium levels. Given this gap in knowledge and the growing interest in how calcium and ROS signaling act to regulate neuronal excitation, we tested whether slight perturbations of ROS produce observable changes in calcium influx and the resultant signaling. First, using cell-specific expression of the genetically encoded, fluorescent calcium indicator GCaMP6f (Akerboom et al., 2013), we were able to visualize calcium transients *in vivo* in the soma of AVA. Using this technique, we observe temporal dynamics (i.e. transient frequency and duration) of calcium transients similar to previous reports (Larsch et al., 2013; Gordus et al., 2015) (Extended Figure 2_1).

To determine if calcium influx is impacted by intracellular concentration of hydrogen peroxide (H_2_O_2_), the most stable and common form of endogenous ROS in cells (Bienert et al., 2006), we subjected wild-type worms expressing GCaMP6f in the AVA to a 5-minute pretreatment of 100 nM H_2_O_2_ prior to imaging. With this technique, we determined that acute treatment with a H_2_O_2_ concentration within the physiological range (10-100 nM; (Sies, 2017)) decreased the total GCaMP activity to 80.9% ± 0.07 of that of untreated controls (n>55 for each group, p=0.006, Figure 2A and 2B, Figure 2_1B). Furthermore, we saw no effect of H_2_O_2_ treatment on baseline fluorescence (Figure 2_1C), indicating that the modest increase in ROS due to this treatment does not detectably change the fluorescence properties of GCaMP itself or basal calcium levels. We conclude that an acute increase in H_2_O_2_ decreases the total calcium influx over the recording time (60 seconds) without drastically modifying the amplitude of calcium influx in the AVA.

**Figure 2:**
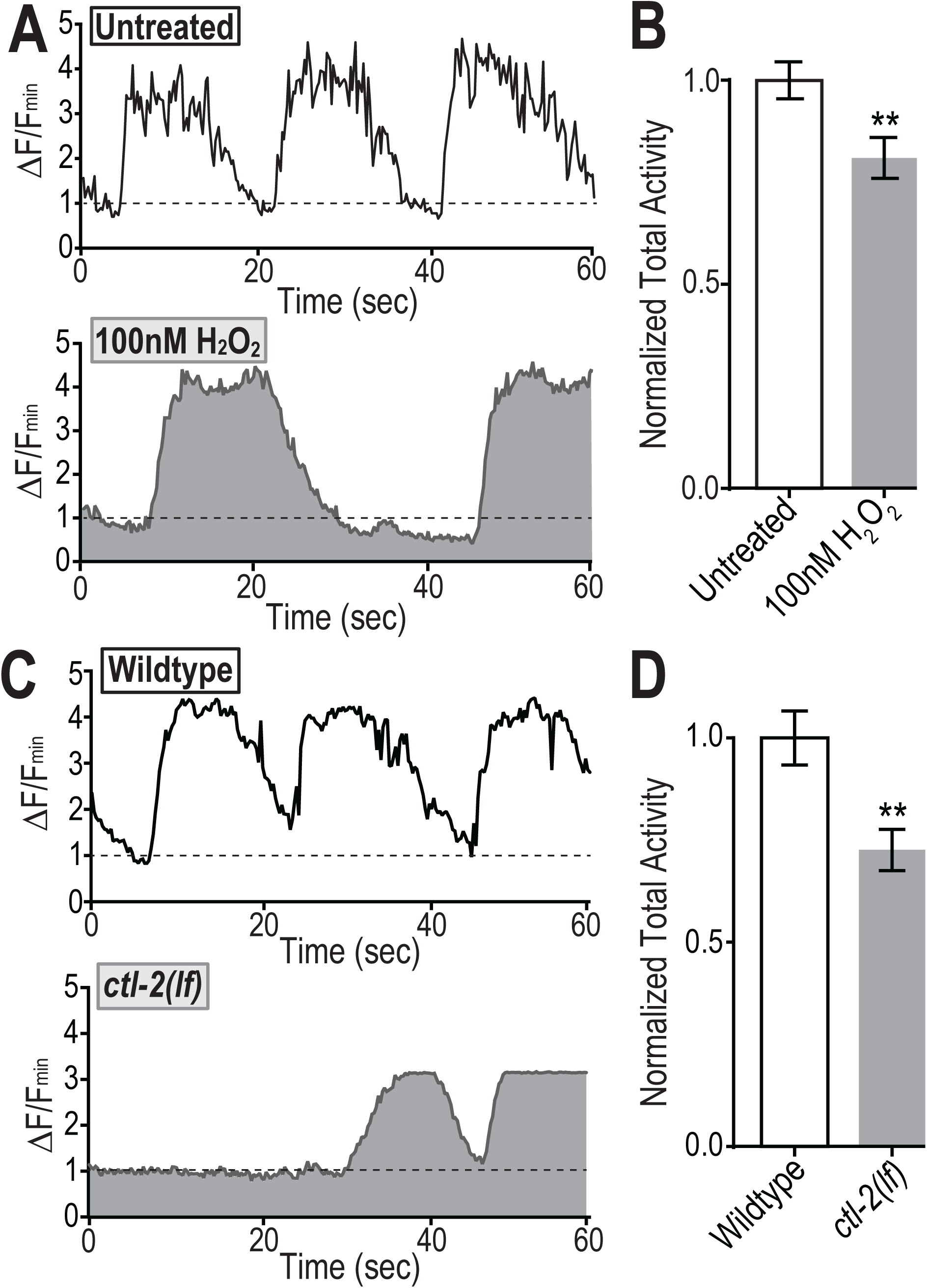
Physiological ROS increase leads to decreased activity-dependent calcium influx. A) Representative traces of somatic GCaMP6f fluorescence in AVA interneurons for 60 seconds, normalized to baseline fluorescence (ΔF/F_min_) from untreated (n=56) and 100 nM H_2_O_2_ treated (n=58) wild-type animals. Dashed line=F_min_. B) Total activity (sum of fluorescence values above baseline normalized to baseline) normalized to untreated wild-type (**: p=0.006). C) Representative traces of somatic GCaMP6f fluorescence in AVA interneurons for 60 seconds, normalized to baseline fluorescence in a wild-type (n=50) and catalase mutants (*ctl-2(lf)*, n=51). Dashed line=F_min_. D) Total activity normalized to wild-type (**: p=0.001).

We sought additional evidence by using a genetic strategy to increase intracellular H_2_O_2_. To this end, we obtained a strain harboring a loss-of-function mutation in the gene encoding the primary *C. elegans* catalase (*ctl-2*), which decomposes approximately 80% of all intracellular H_2_O_2_ to water, including in neuronal tissue (Petriv and Rachubinski, 2004). In these catalase mutants, total GCaMP activity is significantly decreased (27.4% ± 0.08 lower, p=0.001; Figure 2B and 2D). Again, the baseline of GCaMP fluorescence in *ctl-2(lf)* was not different than in wild-type animals (Figure 2_1D). These results support that both acute and chronic increases in H_2_O_2_ within the physiological range decrease the total activity-dependent influx of calcium into *C. elegans* neurons. Based on the identified regulatory role of calcium influx on GLR-1 transport, we then hypothesized that these same modest increases in ROS may affect GLR-1 transport and delivery to synapses.

### Physiological ROS signaling regulates AMPAR transport and delivery to synapses

To test if ROS levels impact GLR-1 transport, we quantified GLR-1 transport as previously described following a 5-minute pretreatment of 10, 50 and 100 nM H_2_O_2_, in which animals swam freely before imaging in the same H_2_O_2_-containing solution. We found that worms treated with H_2_O_2_ had significantly fewer AMPAR transport events ranging from 14.65 ± 1.59 events at 10nM H_2_O_2_ to 11.35 ± 1.14 at 100nM H_2_O_2_ (compared to 23.0 ± 1.09 events in untreated animals, n=22, p<0.0001, Figure 3A and 3B). Given, that all treatments affected GLR-1 transport to a similar degree, we used 50nM of H_2_O_2_ in all following experiments. In addition, quantification of GLR-1 transport in *ctl-2(lf)* mutants revealed a significant decrease in GLR-1 transport events (12.3 ± 1.10 events per kymograph, n=28, p=0.0012; wild-type 19.0 ± 1.71 events, n=20, Figure 3C and 3D). The H_2_O_2_ treatment also led to significantly decreased anterograde transport velocities (1.31 ± 0.02 μm/sec, n=107 events, p<0.0001) compared to untreated controls (1.52 ± 0.03 μm/sec, n= 90 events, Figure 3_1A and C). The *ctl-2(lf)* mutation caused a 17% decrease (1.10 ± 0.02 μm/sec, n=107 events, p<0.0001) compared to wild-type (1.33 ± 0.02 μm/sec, n=59, Figure 3_1D and F). Neither H_2_O_2_ treatment nor *ctl-2(lf)* altered retrograde transport velocity (data not shown), so remaining velocity analyses were focused on that of anterograde transport. In addition, the percent of time GLR-1 containing vesicles spent paused was decreased by these modest elevations in ROS levels (p<0.0001, Figure 3_1B and E). Thus, both acute and chronic increase of H_2_O_2_ lead to decreased transport of GLR-1 containing vesicles to and from the cell body, as would be predicted by our observations of decreased calcium influx in response to modest ROS elevations. Furthermore, these slight increases in ROS were sufficient enough to alter the normal characteristics of vesicle transport including transport velocity as well as the frequency and/or duration of stops along the neuronal process.

**Figure 3:**
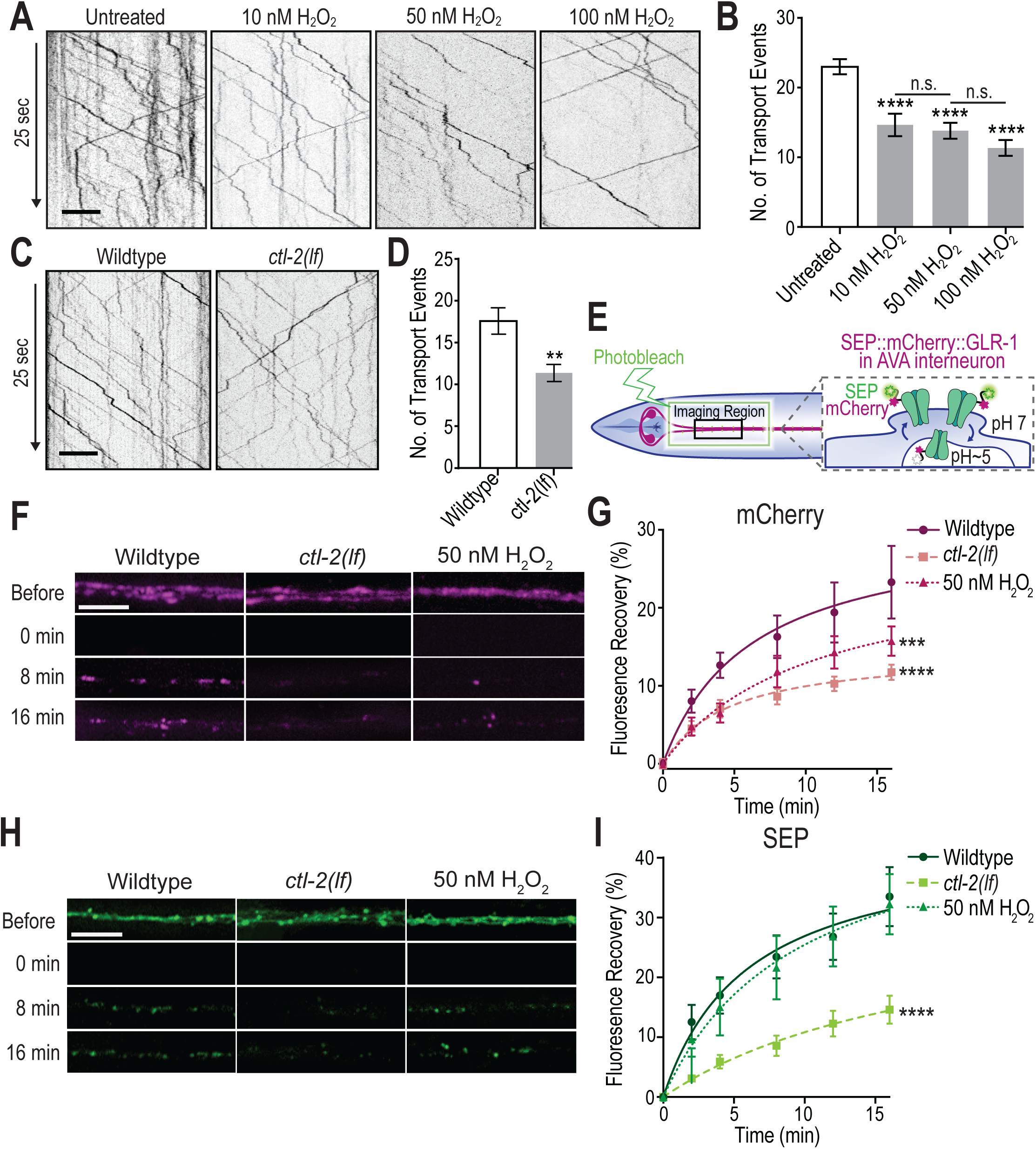
Physiological ROS increase causes decrease in GLR-1 transport and delivery. A) Representative kymographs from untreated, 10, 50 and 100 nM H_2_O_2_ treated wild-type worms. Scale bar=5 μm. B) Quantification of total transport events during a 50 second recording from untreated (n=22), 10 (n=20), 50 (n=21), and 100 nM H_2_O_2_ treated (n=22) wild-type worms (****: p<0.0001, n.s.=not significant). C) Representative kymographs from wild-type and *ctl-2(lf).* D) Quantification of total transport events in wild-type (n=21) and *ctl-2(lf)* (n=28, **: p=0.0012). E) Illustration of imaging location of fluorescence recovery after photobleaching (FRAP) in the AVA interneurons expressing GLR-1 tagged with SEP, a pH-sensitive GFP, and mCherry. F) Representative maximum projections of the mCherry fluorescence in the AVA interneurons before, immediately after (0 min), 8 and 16 minutes after photobleaching of the imaging region. Scale bar = 5 μm. G) Percent recovery of mCherry fluorescence after photobleaching over time in untreated wild-type, untreated *ctl-2(lf)*, and 50 nM H_2_O_2_ treated wild-type worms (n=10 for each group, **: p=0.001, ***: p<0.0001). H) Representative maximum projections of the SEP fluorescence in the AVA interneurons before, immediately after, 8 and 16 minutes after photobleaching of the imaging region. Scale bar = 5 μm. I) Percent recovery of SEP fluorescence compared to before photobleaching from the same worms as in Figure 3F (***: p<0.0001).

Altogether, these changes in GLR-1 transport could impact GLR-1 delivery and exocytosis at synapses. To test this hypothesis, we turned to fluorescence recovery after photobleaching (FRAP) using GLR-1 tagged with mCherry and SEP (a pH-sensitive for of GFP) at the N-terminus (Figure 3E) as previously described (Kennedy et al., 2010; Hoerndli et al., 2013). We monitored recovery of mCherry signal from the dual tagged GLR-1 after bleaching to quantify new delivery of GLR-1 to synaptic sites in the proximal region of the AVA processes. SEP fluorescence, on the other hand, is quenched while in acidic endosomes and therefore protected from photobleaching (Kennedy et al., 2010; Hoerndli et al., 2015), meaning its signal is revealed once released to the membrane providing a measure of exocytosis of GLR-1-containing receptors to synapses (Figure 3E). Worms were again pretreated for 5 minutes with 50 nM H_2_O_2_ then immediately mounted for imaging without any change in solution. During treatment, the rate of GLR-1 delivery was significantly decreased (as determined by the nonlinear fit of the percentage of fluorescence recovery throughout the 16 minutes following photobleaching, n=10, p=0.0011) in comparison to untreated wild-type worms (n=10, Figure 3F and 3G). Similar to the acute H_2_O_2_ treatment, *ctl*-2*(lf)* also led to a significant reduction in the rate of synaptic GLR-1 delivery (n=10, p<0.0001, Figure 3F and 3G). Although identical at 1 and 2 minutes of recovery, *ctl-2(lf)* caused a slightly slower rate of synaptic GLR-1 recovery than acute H_2_O_2_ treatment at later time points. This decrease in delivery could in part be due to the decreased time in which transported vesicles are stopped when ROS is elevated (Figure 3_1B and 3_1E), which likely perturbs the ability to be delivered to synaptic sites. Together, these results show that even a modest, acute elevation of ROS can lead to decreased GLR-1 delivery at synapses.

The efficacy of excitatory neurotransmission is determined in part by the number of AMPARs at the surface of synapses (Huganir and Nicoll, 2013; Henley and Wilkinson, 2016) and although ROS elevation decreased GLR-1 delivery to synapses, it may not affect exocytosis rates or the number of receptors at the synaptic membrane. To determine if this is the case, we quantified the SEP signal following GLR-1 photobleaching in both acute and genetic conditions of ROS elevation. Interestingly, the exocytosis rate of GLR-1 seemed to be unaffected by acute H_2_O_2_ treatment whereas *ctl-2(lf)* significantly decreased GLR-1 exocytosis rates (n=10 for all groups, p<0.0001, Figure 3H and 3I). This difference in SEP recovery suggests that acute and chronic ROS elevation differentially affect GLR-1 exocytosis at synapses. Chronic ROS elevation could lead to sustained decreases in GLR-1 delivery resulting in a time-dependent depletion of the synaptic reserves required for the GLR-1 exocytosis. If true, then a marked decrease in SEP::mCherry::GLR-1 signal at steady state in *ctl-2(lf)* mutants would be expected. Surprisingly, measurements of SEP::mCherry::GLR-1 fluorescence along the neuronal process before FRAP in *ctl-2(lf)* and wild-type animals did not show a significant change in mCherry or SEP signal (data not shown). We postulated that the overexpression of GLR-1 necessary to follow single vesicle dynamics, might mask changes in steady state.

To quantify the effect of physiological elevation of ROS levels in *ctl-2(lf)* mutants on global glutamatergic circuit function and circumvent potential overexpression issues of the SEP::mCherry::GLR-1 in AVA, we turned to behavioral analysis. The spontaneous reversal of *C. elegans* animals has been shown to reflect the function and number of synaptic GLR-1 receptors (Zheng et al., 1999; Burbea et al., 2002; Park et al., 2009; Monteiro et al., 2012). In addition, AVA activation has been shown to be essential for spontaneous reversals (Gray et al., 2005; Ben Arous et al., 2010). Thus, we obtained reversal data for *ctl-2(lf)* and found that *ctl-2(lf)* mutants exhibited fewer spontaneous reversals (1.37 ± 0.19 reversals per minute, p<0.0001, n=38) compared to wild-type (3.78 ± 0.26, n=37, data not shown). This behavioral change supports our calcium imaging data in which the total activity of AVA is decreased in *ctl-2(lf)*. Altogether, our results clearly show that modest ROS elevation is sufficient to modify synaptic GLR-1 transport, delivery and, with chronic elevations, exocytosis to synapses ultimately affecting glutamatergic circuit function.

### Increased ROS modulate GLR-1 transport at or directly downstream of L-type VGCC

Our results indicate that modest increases in ROS (within the physiological range for neurons; Sies, 2017) decrease activity-dependent calcium influx in the AVA interneurons of *C. elegans* (Figure 2). This results in decreased GLR-1 transport into and out of the cell body as well as decreased in anterograde transport velocity and time vesicles spent stopped along the neurite (Figure 3A-D, Figure 3_1). Together, these changes in transport likely contribute to decreased delivery and, in the case of prolonged elevations in ROS, exocytosis of GLR-1 at synapses (Figure 3E-I). To add clarity, we investigated mechanisms by which excess ROS decreases calcium influx, CaMKII activation, and therefore GLR-1 transport.

Our data presented above suggest a regulatory signaling pathway (Figure 4A), in which depolarization of AVA via glutamate receptor activation leads to calcium influx through VGCCs to activate CaMKII. This model led us hypothesize that elevated ROS could be affecting calcium signaling needed for CaMKII activation. We used a genetic epistasis strategy to test whether this is a linear pathway in which ROS function upstream of CaMKII activation to modulate GLR-1 transport. We made double mutants with *unc-43(lf)* and *ctl-2(lf)* as well as *unc-43 (gf)* and *ctl-2(lf).* If ROS are acting upstream of CaMKII activation, then one would predict that *unc-43(gf/lf); ctl-2(lf)* double mutants would be similar to the single *unc-43* mutations alone as the modest decrease in calcium due to *ctl-2(lf)* would not affect the activation of calcium-independent CaMKII in the *unc-43(gf)* mutants.

**Figure 4:**
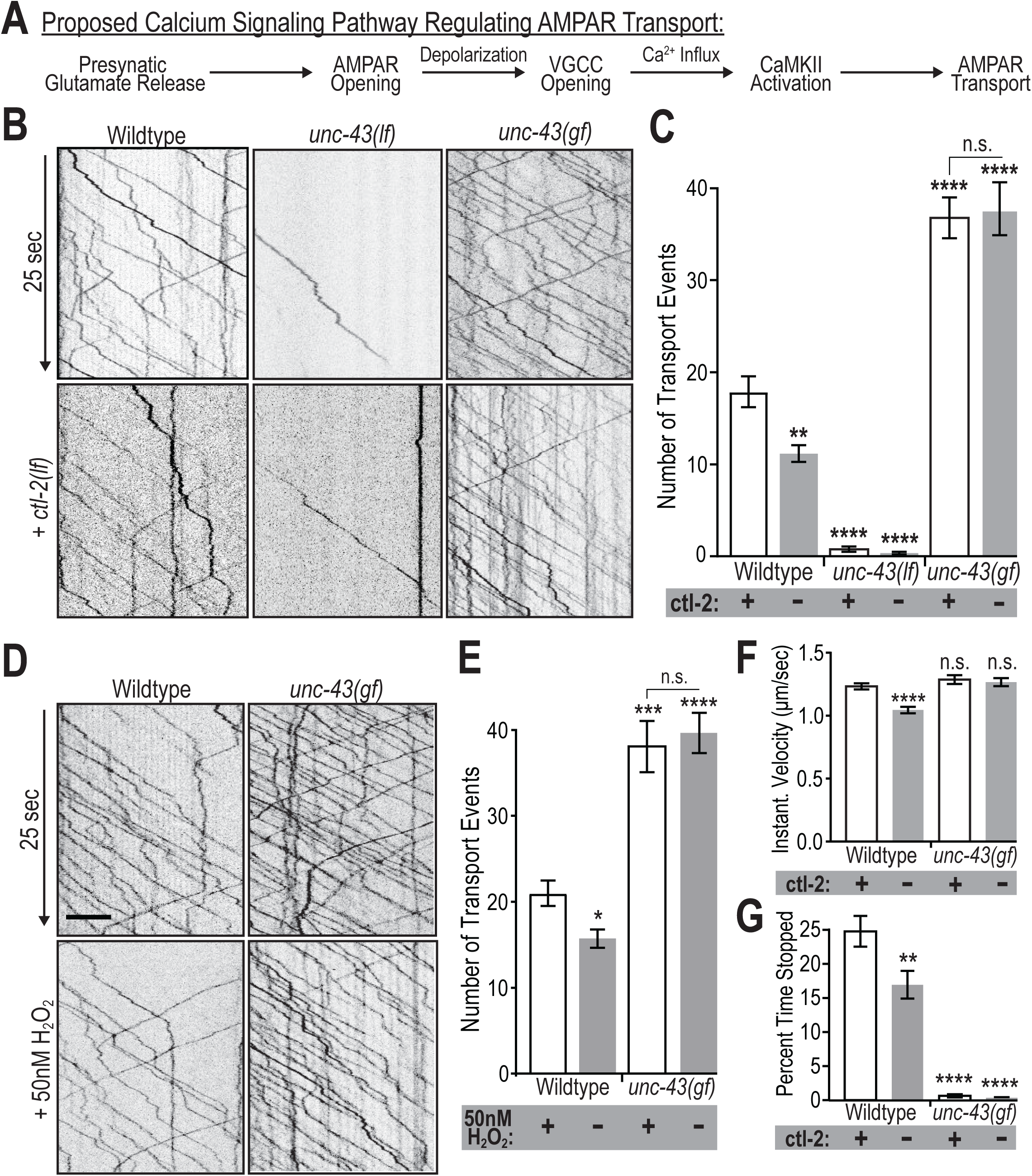
Increased ROS acts on GLR-1 transport upstream of CaMKII activation. A) Previously proposed model of an activity-dependent calcium signaling pathway regulating AMPAR transport (Hoerndli et al., 2015). B and D) 25 seconds of representative kymographs from each experimental group. Scale bar=5 μm. C) Quantification of transport events from full-length kymographs (50 s) in wild-type, *unc-43*(lf), *unc-43*(gf) with (+, white bars) and without (-, grey bars) the *ctl-2* gene (n≥13; n.s.=not significant, **: p=0.0012, ****: p<0.0001 compared to wild-type containing *ctl-2*). E) Quantification of transport events in wild-type or *unc-43(gf)* with (+, grey bars) or without (-, white bars) 50 nM H_2_O_2_ treatment (n≥14; *: p=0.025, ***: p=0.0002, ****: p<0.0001 compared to untreated). F) Instantaneous velocity of anterograde transport for the same animals from part B and C (n>60 events, ****: p<0.0001 compared to wild-type animals with *ctl-2*). G) Percent of time GLR-1 vesicles spent stopped in these same genotypes (n>60 events, **: p=0.009, ****: p<0.0001 compared to wild-type).

To test this, we analyzed mCherry::GLR-1 transport and again observed that *ctl-2(lf)* alone decreases GLR-1 transport, but interestingly, the addition of the *ctl-2(lf)* to *unc-43 (lf or gf)* mutations does not change the amount of transport compared to *unc-43* mutation alone (Figure 4B and C). Analysis of anterograde transport velocity and stopping in the *unc-43(gf)* mutants mirror these results in that the velocity and stopping is the same in the *unc-43(gf); ctl-2(lf)* as in the *unc-43(gf)* single mutant (Figure 4F and G). It is interesting to note that the addition of the *ctl-2(lf)* mutation to *unc-43(gf)* did not affect transport velocity when compared with *unc-43(gf)*, which indicates that modest ROS elevation likely does not lead to a global perturbations of cellular processes which would indirectly affect molecular motor-dependent activity. To determine whether this is unique to *ctl-2(lf)*, we also treated *unc-43(gf)* animals with 50 nM H_2_O_2_ prior to imaging transport. Similar to *ctl-2(lf)*, acute treatment with H_2_O_2_ had no effect on the *unc-43(gf)* mediated increase in GLR-1 transport (Figure 4D and 4E). Together, these data indicate that ROS signaling regulates GLR-1 transport upstream of CaMKII activation.

To pinpoint potential targets of ROS upstream of CaMKII activation, we next investigated L-VGCCs, which we have shown to play an important role in regulating GLR-1 transport events and delivery (Figure 1 and 3). To determine if L-VGCCs or downstream calcium signaling is a target of ROS modulation, we used a genetic epistasis approach. We made double mutants with *ctl-2(lf)* and either *egl-19(rf)* or *egl-19(gf)*, then analyzed GLR-1 transport. These analyses reveal that the amount of GLR-1 transport in *egl-19(rf); ctl-2(lf)* is not significantly different from *egl-19(rf)* (Figure 5A and 5B). However, *egl-19(gf); ctl-2(lf)* mutants have significantly decreased GLR-1 transport events in comparison to *egl-19(gf)* alone (13.9 ± 1.17 vs 26.7 ±1.76 respectively, n=16, p<0.0001, Figure 5A and 5B). In fact, the double mutant was not significantly different than *ctl-2(lf)* alone (11.2 ± 0.91 events, n=19, p=0.42, Figure 5A and 5B). Quantification of the anterograde velocity and stopping of transport events in these strains further support the idea that ROS affect calcium signaling at or just downstream of L-VGCCs. In the *egl-19(gf)* mutant, instantaneous velocity of anterograde transport is increased compared to wild-type (1.37 ± 0.03 vs 1.23 ± 0.02 μm/sec, n>40 events, p=0.0005, Figure 5_1A). When combined with *ctl-2(lf)*, anterograde transport velocity is decreased to around that of the *ctl-2(lf)* mutation alone (1.12 ± 0.03, n=43, vs 1.04 ± 0.02, n=82, p=0.66, Figure 5_1A and D). The percent of time GLR-1 vesicles spent stopped along the neuronal process was increased in the *egl-19(gf)* mutants (stopped 29.9% ± 2.72 of time, n=96) compared to wild-type (20.8% ± 1.86, n=95, p=0.044). When combined with *ctl-2(lf)*, stopping was drastically decreased compared to *egl-19(gf)* alone (11.65% ± 1.94, n=44, p<0.0001) and wild-type (20.88% ± 1.86, n=89, p=0.015, Figure 5_1B). These data suggest that elevated ROS change transport velocity and stopping throughout the neurite via decreased calcium influx. If true, then we hypothesized that transport velocity and stopping should be decreased in the *egl-19(rf)* mutant.

**Figure 5:**
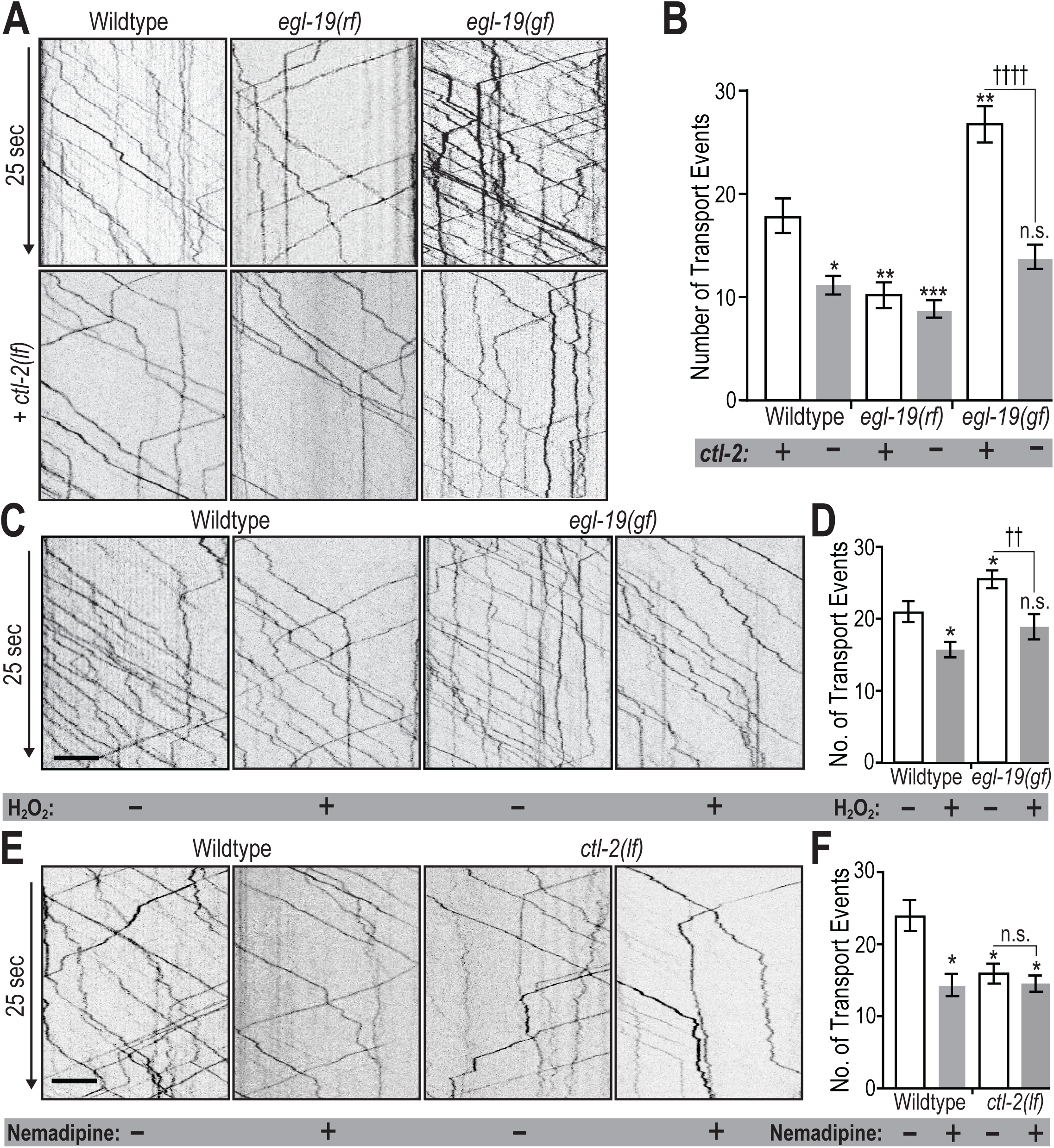
ROS decrease GLR-1 transport by acting on or downstream of L-VGCC. A, C and E) 25 seconds from representative kymographs from each experimental group. Scale bar=5 μm. B) Quantification of transport events from wild-type, *egl-19*(rf), or *egl-19(gf)* with (+, white bars) or without (-, grey bars) the *ctl-2* gene (n≥15; n.s.=not significant, *: p=0.012,**: p=0.007, ***: p=0.0005, compared to wild-type with the *ctl-2* gene; ††††: p<0.0001 compared to *egl-19(gf)* alone). D) Quantification of transport events from wild-type and *egl-19(gf)* without (-, white bars) or with (+, grey bars) 50 nM H_2_O_2_ treatment (n≥18; *: p≤0.038 compared to untreated wild-type; ††: p=0.007 compared to untreated *egl-19(gf)*). F) Quantification of transport events in wild-type and ctl-2(lf) with (+, grey bars) and without (-, white bars) 10 μM nemadipine treatment (n>10; *: p<0.05 compared to untreated wild-type).

Similar to our quantification of GLR-1 transport numbers, the transport velocity of *egl-19(rf)*, is decreased compared to wild-type (1.08 ± 0.04 μm/sec, n=86, p=0.0001) and is unaffected by the addition of the *ctl-2(lf)* mutation (Figure 5_1A and C). The percentage of time vesicles were stopped was also decreased in *egl-19(rf)* mutants (14.93% ± 2.15, n=79, p=0.001). However, in contrast to our velocity results, we observed a slight, but not significant decrease in stopping with the addition of *ctl-2(lf)* (8.7% ± 1.60, n=67, p=0.43, Figure 5_1B). These data indicate that the *ctl-2(lf)* mutation is acting in the same pathway as *egl-19* to regulate GLR-1 transport. To ensure that these results are due to elevated ROS levels and not developmental effects of *ctl-2(lf)*, we also treated *egl-19(gf)* animals with 50 nM H_2_O_2._ As previously observed, H_2_O_2_ treatment of wild-type animals reduced GLR-1 transport and reduced *egl-19(gf)* transport to the same level as H_2_O_2_ alone (26.11 ± 1.15 vs 18.9 ± 1.76 events, n>18, Figure 5C and 5D). To eliminate the possibility of these results being caused by developmental changes in the *egl-19(gf/lf)* mutants, we also analyzed GLR-1 transport in wild-type and *ctl-2(lf)* mutants following treatment with the L-VGCC blocker, nemadipine. As we previously showed, a 10 μM nemadipine treatment significantly reduced the total amount of GLR-1 transport (14.36 ± 1.53 events, n=11, compared to controls 24.00 ± 2.15 events, n=10; p=0.008). However, nemadipine treatment did not further reduce transport in *ctl-2(lf)* mutants (15.92 ± 1.38 events, n=13, in untreated compared to 14.56 ± 1.13 events with nemadipine, n=9, Figure 5E and 5F). Altogether, these observations suggest that ROS elevation is affecting L-VGCC/EGL-19 dependent calcium signaling.

To obtain additional insight into how ROS modulate GLR-1 transport and delivery, we combined FRAP of SEP::mCherry::GLR-1 and our genetic epistasis strategy (Figure 6). We quantified synaptic delivery of GLR-1 using FRAP of mCherry signal (Figure 6A and 6B) and synaptic exocytosis using FRAP of SEP signal (Figure 6C and 6D) in single *egl-19(rf)* or *egl-19(gf)* mutants and when doubled with *ctl-2(lf)*. Similar to our GLR-1 transport results, quantification of mCherry and SEP FRAP showed that the rate of GLR-1 delivery and exocytosis is decreased in *egl-19(rf)* mutants and addition of *ctl-2(lf)* causes no change in those rates (n=10, Figure 6). Alternatively, the FRAP of mCherry in *egl-19(gf); ctl-2(lf)* is reduced compared to *egl-19(gf)*, but not identical to *ctl-2(lf)* (Figure 6B) as is the case with the FRAP of SEP in these genotypes (Figure 6D). These results suggest that ROS not only modulate GLR-1 transport, but also delivery and exocytosis by affecting the signaling cascade initiated by calcium influx through L-VGCC/EGL-19. Future work will determine if elevated ROS changes calcium influx by directly modulating L-VGCCs or by acting on downstream components such as calmodulin and CaMKII.

**Figure 6:**
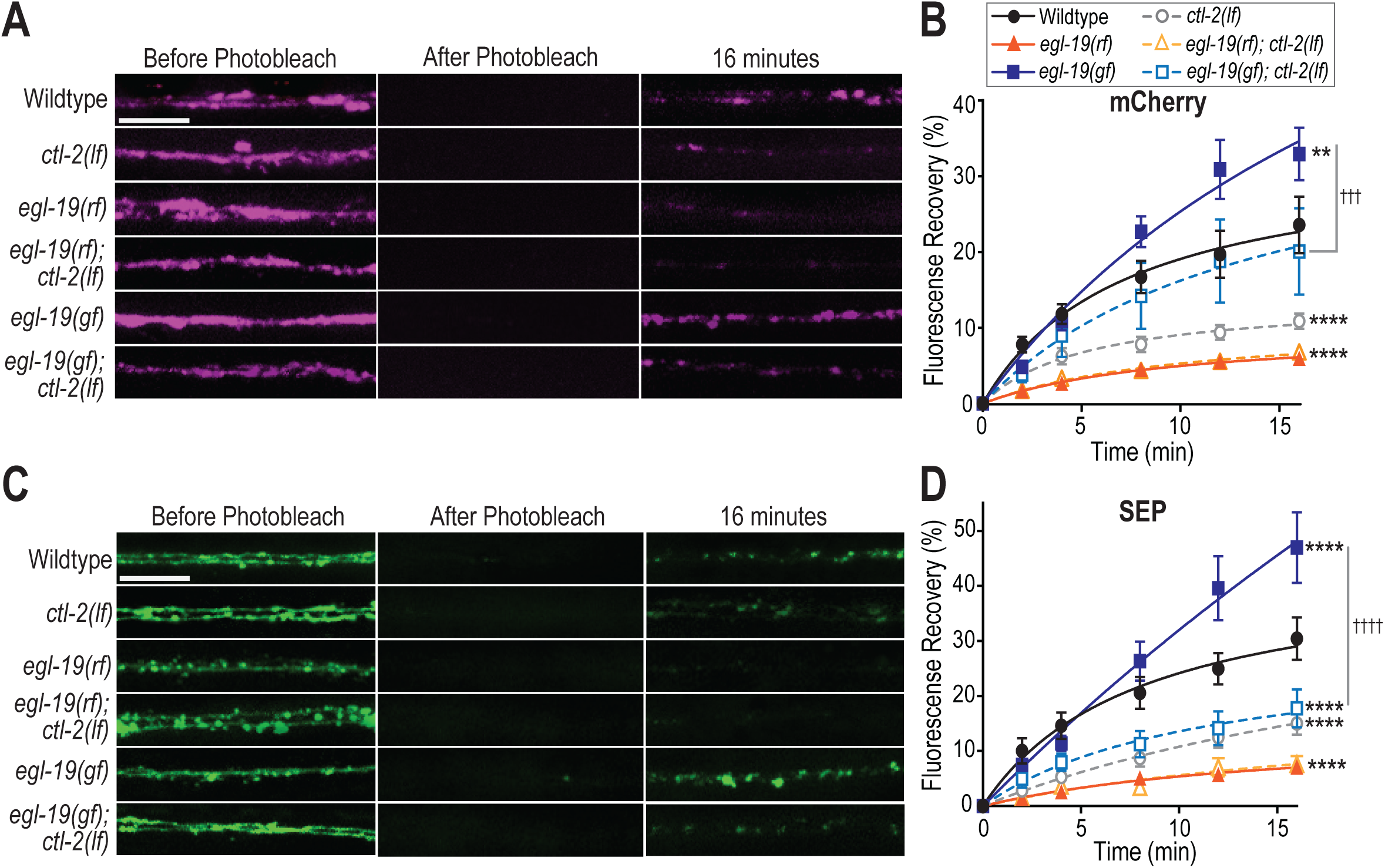
ROS decrease GLR-1 synaptic delivery and exocytosis by acting on or downstream of L-VGCC. A) Representative maximum projections of the mCherry fluorescence in the AVA interneurons before, immediately after and 16 minutes after photobleaching of the imaging region from each experimental group. Scale bar=5 μm. B) Recovery of mCherry fluorescence following photobleaching in wild-type, *egl-19(lf)* and *egl-19(gf)* with (dashed curves) and without (solid curves) *ctl-2(lf)* (n≥8; **: p=0.003, ****: p<0.0001 compared to wild-type; †††: p=0.0001 compared to *egl-19*(gf) single mutant). C) Representative maximum projections of the SEP fluorescence in the same groups. Scale bar=5 μm. D) SEP fluorescence recovery in the same experimental groups (****: p<0.0001 compared to wild-type; ††††: p<0.0001 compared to *egl-19(gf)*).

Our data provide a better understanding of the mechanism by which calcium influx regulates AMPAR transport to ultimately affect synaptic delivery and exocytosis of AMPARs. We go on to show that calcium influx itself is modified by ROS. This leads to our suggested model (Figure 7) in which physiological elevations of ROS can modulate synaptic GLR-1 transport at the soma and in the dendritic process by modifying calcium influx, originating to a large extent from L-VGCCs. Taken together, our results identify a novel role for oxidant signaling in the regulation of synaptic AMPAR transport and delivery, which in turn could be critical for coupling the metabolic demands of neuronal activity with excitatory neurotransmission. To test this model and understand its biological significance, future studies are required. In particular, it will be critical to know if and how ROS production is modulated by neuronal activity.

**Figure 7:**
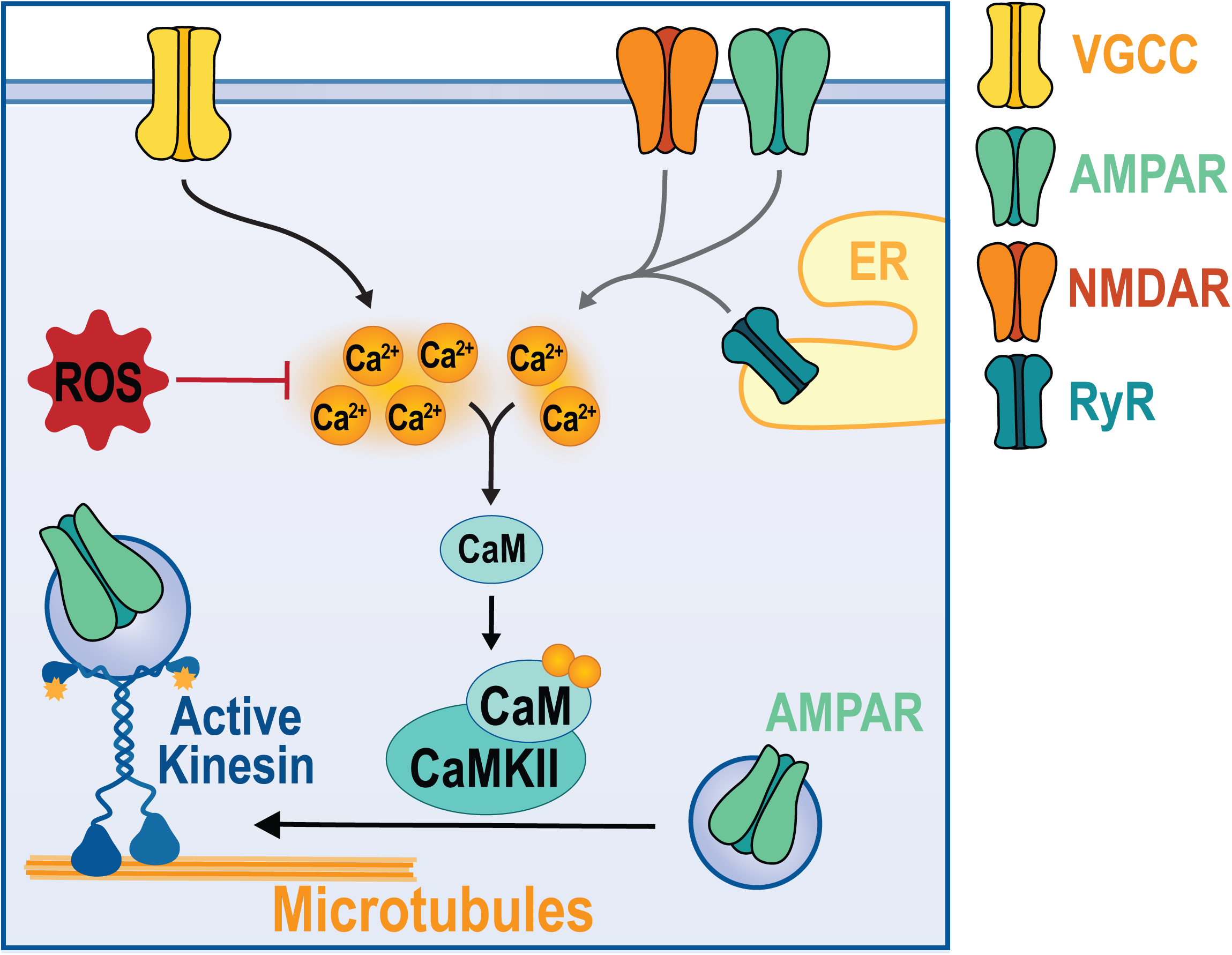
Proposed ROS signaling effect on GLR-1 transport. Model showing that increases in ROS inhibit the activity-dependent calcium influx, likely from L-VGCC (yellow channel, VGCC). As a consequence, this limits calmodulin (CaM) binding and activation of CaMKII reducing the number of AMPAR (green channel) transport and delivery to synapses.

## DISCUSSION

Activity-dependent calcium signaling is required for AMPAR transport in vertebrate and *C. elegans* glutamatergic neurons (Hoerndli et al., 2015; Hangen et al., 2018). Here we show that in *C. elegans*, calcium influx is not only required, but that the magnitude of calcium influx directly corresponds to changes in amount of AMPAR transport to and from the cell body as well as AMPAR transport dynamics within neuronal processes. Furthermore, we show that physiological increases in ROS decrease somatic calcium influx (Figure 2) causing a subsequent decrease in somatic AMPAR export and synaptic delivery (Figure 3). This study reveals a physiological interplay between ROS and calcium signaling that directly affects AMPAR transport and delivery (Figure 7). It also prompts fundamental questions regarding how AMPAR transport is regulated by activity in a spatially specific manner, such as how do somatic vs local dendritic calcium signaling pathways regulate AMPAR transport and what are their effects on synaptic function and plasticity?

Long-distance AMPAR transport is one of the least characterized components of the multistep trafficking of AMPAR from the cell body to the periphery (Henley and Wilkinson, 2016). It is clear that transport requires molecular motors, including kinesin-1 (Kim and Lisman, 2001; Setou et al., 2002; Hoerndli et al., 2013), and is regulated by neuronal activity and CaMKII (Esteves da Silva et al., 2015; Hoerndli et al., 2015; Hangen et al., 2018). However, we do not understand the roles of other key players required for activity-dependent regulation and at which steps (i.e. cargo loading, transport, delivery and removal) they act. Consistent with previous reports (Hoerndli et al., 2015), we show decreased somatic export of AMPARs in mutants with reduced L-VGCC function or after treatment with the L-VGCC blocker, nemadipine. On the contrary, increased calcium influx in gain-of-function L-VGCC mutants leads to more AMPAR transport demonstrating that activity-dependent calcium influx through L-VGCCs bidirectionally regulate AMPAR transport out of the cell body.

Once motors and cargoes have entered the dendrites, they are subject to the conditions of those compartments. For instance, GLR-1 transport in the AVA neurite is bidirectional and heterogenous, displaying increased stopping probabilities at synaptic localizations of GLR-1 (Hoerndli et al., 2013). However, it was unclear what factors in the neurite might regulate stopping probability. Here, we show how manipulations of calcium influx through L-VGCC modify AMPAR transport dynamics, such as stops, velocity profiles (Figure 1_1) and synaptic delivery (Figure 6). We show that decreased calcium influx through L-VGCCs decreases anterograde velocity and stopping of AMPAR-containing vesicles within the neurite. Conversely, increasing calcium influx through L-VGCC led to increased anterograde velocities and pausing (Figure 1_1).

Decreased velocity and stopping corresponds to less GLR-1 delivery and insertion to synapses whereas higher velocities and stops correspond to higher delivery and insertion (Figure 6F). These findings concur with studies using vertebrate neurons in which long-term increases in dendritic calcium increased the amount of AMPAR transport and delivery, but contradicts observations following acute calcium depletion (Hangen et al., 2018). Nevertheless, our data is a new in vivo demonstration of activity-dependent calcium signaling bidirectionally regulating AMPAR transport through the neuron determining transport vesicle numbers, dynamics and delivery at synapses at short and long timescales.

Importantly, gain-or loss-of-function CaMKII mutants have a much stronger effect on GLR-1 somatic export than L-VGCC mutations alone. This could be due to the mutations in L-VGCCs not being a complete loss-or gain-of-function. However, it could also suggest that additional signaling pathways converge onto CaMKII to regulate GLR-1 somatic export. These signaling pathways could involve other calcium sources (i.e. NMDAR and ryanodine receptors) or other signaling cascades such as those involving protein kinase A/C or mitogen-activated protein kinase (Boehm et al., 2006; Zheng and Keifer, 2008; Ren et al., 2013; Eales et al., 2014; Tang and Yasuda, 2017). Although these signaling pathways change synaptic AMPAR trafficking, their role in somatic export and/or long-distance transport dynamics is unknown.

### Compartmentalized calcium signaling

Calcium influx into neurons at the soma, dendrites and axons differentially regulates a variety of neuronal functions (reviewed in Brini et al., 2014). Calcium influx in the dendritic shaft and dendritic spines comes from various ion channels including AMPAR, NMDAR and voltage-sensitive calcium channels localized to the membrane, endoplasmic reticulum and mitochondria (Catterall, 2011; Higley and Sabatini, 2012). Calcium imaging in cell culture has shown calcium to remain within the spine following single spine activation (Sabatini et al., 2001; Nimchinsky et al., 2002). Interestingly, vertebrate and invertebrate neurons have compartmentalized calcium transients that are important for neuronal function and computation (Higley and Sabatini, 2012; Ali and Kwan, 2019; Donato et al., 2019). However, it is unclear how compartmentalization of calcium regulates downstream processes such as trafficking of synaptic proteins.

Compartmentalized calcium signaling is conserved in *C. elegans* interneurons (Hendricks et al., 2012; Donato et al., 2019). Our data shows that AMPAR transport in the long process of AVA is affected by altered calcium influx due to mutations in L-VGCC and ROS levels. Some of these effects are likely to be local in scope. However, we have no knowledge regarding distribution of VGCCs, including EGL-19, or about compartmentalization of calcium signaling in the AVA. It is unclear how synaptic inputs and calcium influx are integrated at dendritic and cellular levels to tailor AMPAR distribution. Our study provides a platform to start understanding the *in vivo* mechanisms regulating AMPAR distribution ultimately affecting synaptic strength and behavior of animals.

### ROS

Neuronal excitation and synaptic plasticity have high energy demands and correlate with physiological fluctuations in ROS levels (Bindokas et al., 1996). Pathophysiological ROS elevation is observed in aging and neurodegenerative conditions such as Parkinson’s, Huntington’s and Alzheimer’s (Stefanatos and Sanz, 2018). Intriguingly, some amount of ROS, in particular superoxide (O_2-_), is required for LTP induction and memory formation (Thiels et al., 2000). This suggests that ROS are required for LTP formation, supporting a necessary signaling role in excitatory neuronal function, perhaps specifically in AMPAR trafficking. However, there is currently no direct mechanistic link between ROS signaling and regulation of AMPAR trafficking. Our results show that physiological ROS elevation decreases AMPAR transport, delivery and exocytosis (Figure 3) through a mechanism involving decreased calcium influx (Figure 2). Consistent with our results, hypoxic conditions triggering increases in ROS production lead to decreased GLR-1 trafficking in *C. elegans* (Park and Rongo, 2016).

Although there have been quite a few studies of the regulation of L-VGCC function by physiological ROS in several cell types (Chaplin and Amberg, 2012; Todorovic and Jevtovic-Todorovic, 2014; Cserne Szappanos et al., 2017) there have been extremely few in neurons (Massaad and Klann, 2010; Hidalgo and Arias-Cavieres, 2016; Wilson et al., 2018) and none *in vivo*. Our study is the first to show how small increases in ROS affect whole cell activity-dependent calcium influx and direct downstream targets such as synaptic AMPA receptor transport and delivery. We show that genetically and pharmacologically induced increases in ROS at physiological signaling levels (∼50 nM; Sies, 2017) leads to decreased calcium influx (Figure 2) in the cell bodies of *C. elegans* AVA neurons. This contrasts what has been found in vertebrate arterial smooth muscle and gonadotropes (Chaplin and Amberg, 2012; Dang et al., 2018), but is consistent with reports of mitochondrial ROS elevation inhibiting L-VGCCs in cardiomyocytes (Scragg et al., 2008). Discrepancies in the concentration of ROS treatment between these studies and the one presented here could explain the opposing results. Several cellular processes have been shown to be impacted differently by ROS levels within vs outside of the physiological range (Wilson and González-Billault, 2015; Beckhauser et al., 2016). It is possible that ROS regulation of calcium sources follows suit in that moderate increases in ROS cause decreased calcium influx but greater increases in ROS cause increased calcium influx.

Overall, our study shows that calcium influx through L-VGCCs bidirectionally regulates somatic export and local dendritic dynamics of AMPAR transport. In addition, it provides the first mechanistic insight on how physiological ROS signaling modulates AMPAR transport with consequences for synaptic delivery and exocytosis. Many unanswered questions remain. Such as, does neuronal activity and calcium trigger ROS signaling? If yes, from where does it originate? What are the temporal and spatial profiles of activity-dependent ROS production? What are the differences between somatic and post-synaptic calcium or ROS signaling for AMPAR transport and synaptic plasticity? The ability to answer these questions will rely on the development of high affinity, genetically encoded ROS sensors enabling *in vivo* subcellular imaging of these dynamics.

## Acknowledgements

Funding for this work was provided by the College of Veterinary Medicine and Biomedical Sciences and the Molecular, Cellular and Integrative Neuroscience Program at Colorado State University (CSU) for Dr. Hoerndli. Dr. Amberg, is supported by R01: HD087347. This work was made possible by the research support from Dr. Bruce Pulford (CSU) and some strains provided by the CGC, which is funded by NIH Office of Research Infrastructure Programs (P40 OD010440). A special thanks to Dr. Attila Stetak (University of Basel, Switzerland) for providing GCaMP6f *C. elegans* reagents.

## FIGURE LEGENDS

**Figure 1_1:**
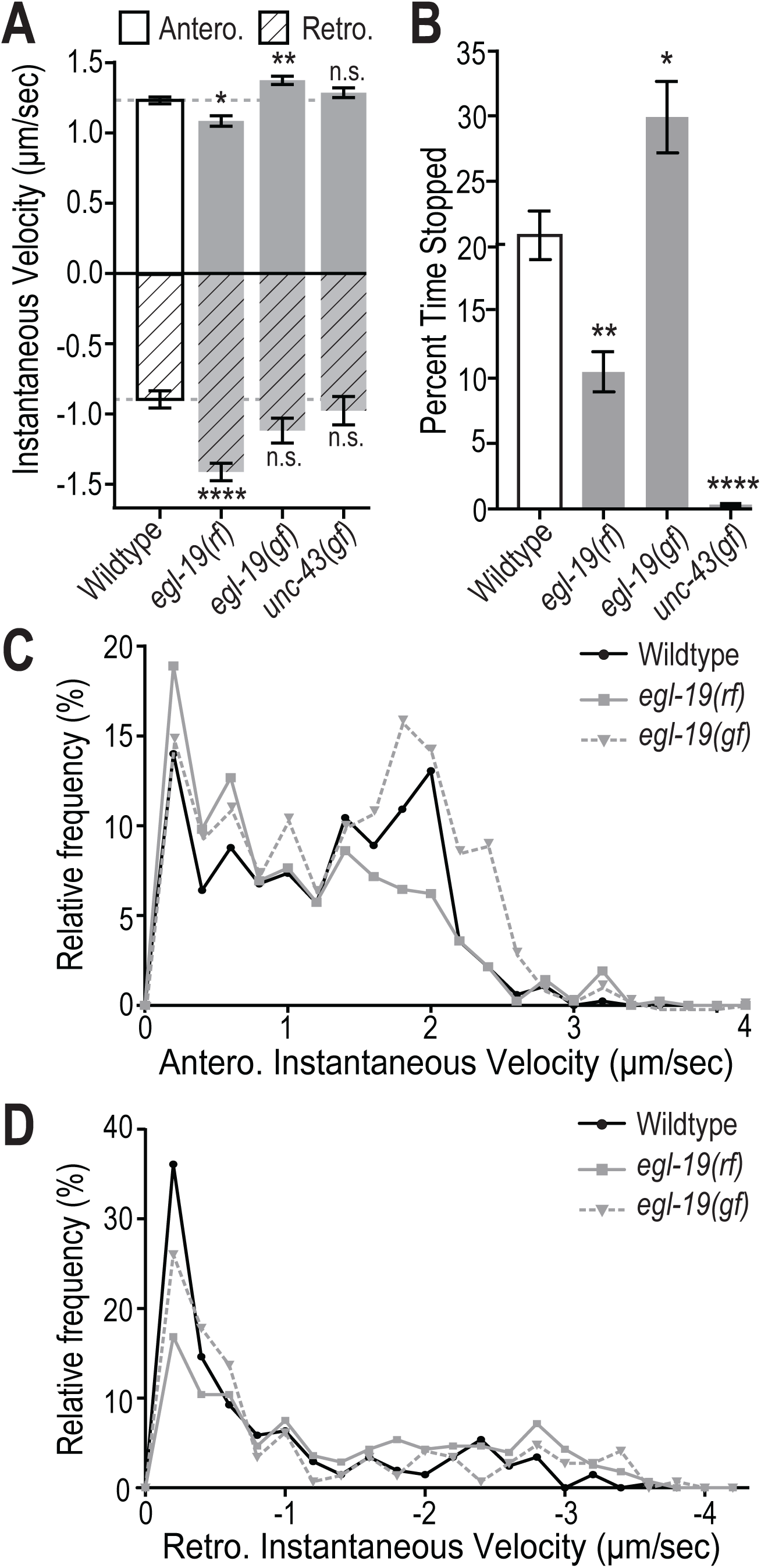
Activity-dependent calcium influx modulates transport velocity and percent of time GLR-1 vesicles spent stopped. A-D) Stop and velocity analysis of more than 60 transport events from wild-type, *egl-19(rf), egl-19(gf)* and *unc-43(gf)* mutants. A) Instantaneous velocity of AMPAR vesicles traveling in either an anterograde (solid) or retrograde fashion (diagonal lines). n>60 transport events, n.s.=not significant, *: p=0.012, **: p=0.003, ****: p<0.0001 compared to wild-type. B) The percent of time GLR-1 vesicles spent stopped in each of the genotypes as quantified from the same transport events in part A. *: p=0.043, **: p=0.001, ****: p<0.0001 compared to wild-type. C and D) Distribution of instantaneous velocities (binned every 0.2 μm/sec) for anterograde (C) and retrograde (D) transport for wild-type, *egl-19(rf)* and *egl-19(gf).*

**Figure 2_1:**
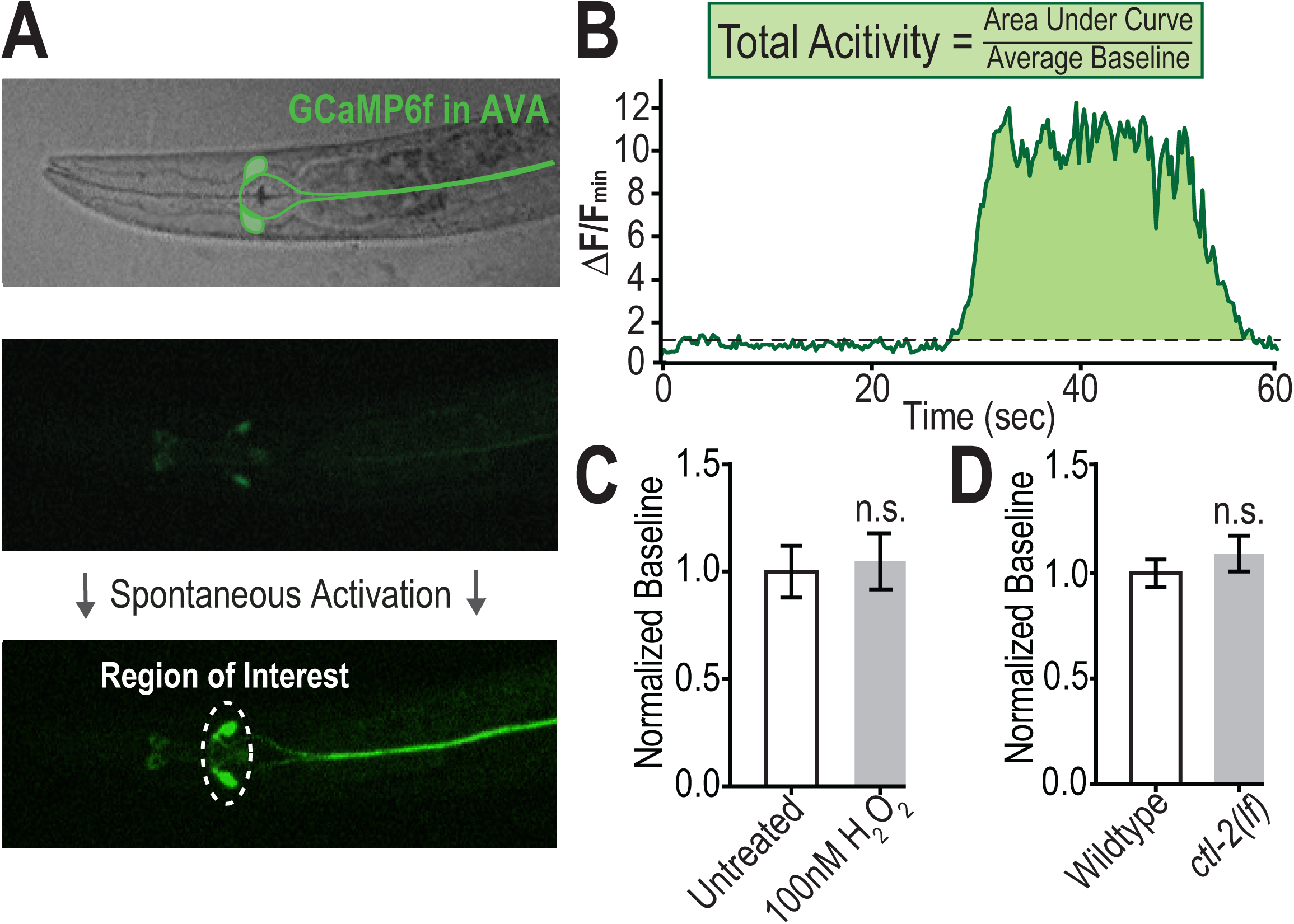
Cell specific expression of GCaMP6f in AVA interneuron. A) Confocal images of the ventral side of a *C. elegans* expressing GCaMP6f in the AVA interneuron. B) GCaMP6f ΔF/F_min_ over time. Grey dashed line represents baseline threshold (30% of minimum fluorescence value). Fluorescence values above that threshold are summed (area under curve) and normalized to the average baseline value to determine the total activity during the 60 second recording. C and D) Average fluorescence baseline of controls, H_2_O_2_ treated and catalase mutants from Figure 2 normalized to corresponding control group (n≥50, n.s. = not significant).

**Figure 3_1:**
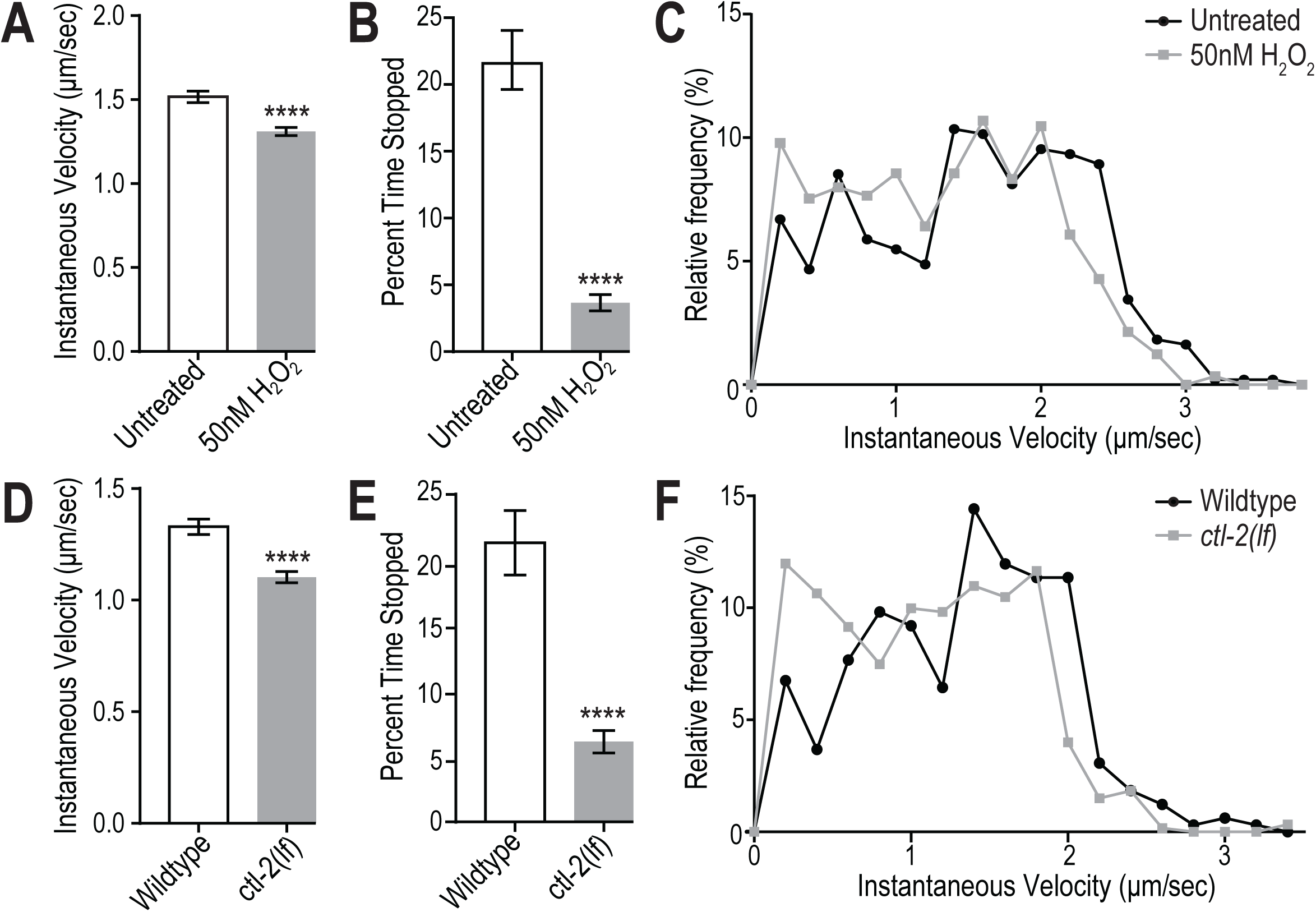
Increased ROS levels decrease velocity and stopping of GLR-1 transport vesicles. A-C) Stop and velocity analysis of anterograde transport events from untreated and 50 nM H_2_O_2_ treated wild-type worms (n>90 events). D-F) Stop and velocity analysis of transport events from wild-type and catalase mutants (*ctl-2(lf)*) (n>50 events). A and D) Instantaneous velocity of anterograde transport in each group, ****: p<0.0001 compared to wild-type. B and E) Percent of time GLR-1 vesicles spent stopped in each group, ****: p<0.0001 compared to wild-type. C and F) Frequency distribution of instantaneous velocities of anterograde transport events within each group (velocities binned every 0.2 μm/sec).

**Figure 5_1:**
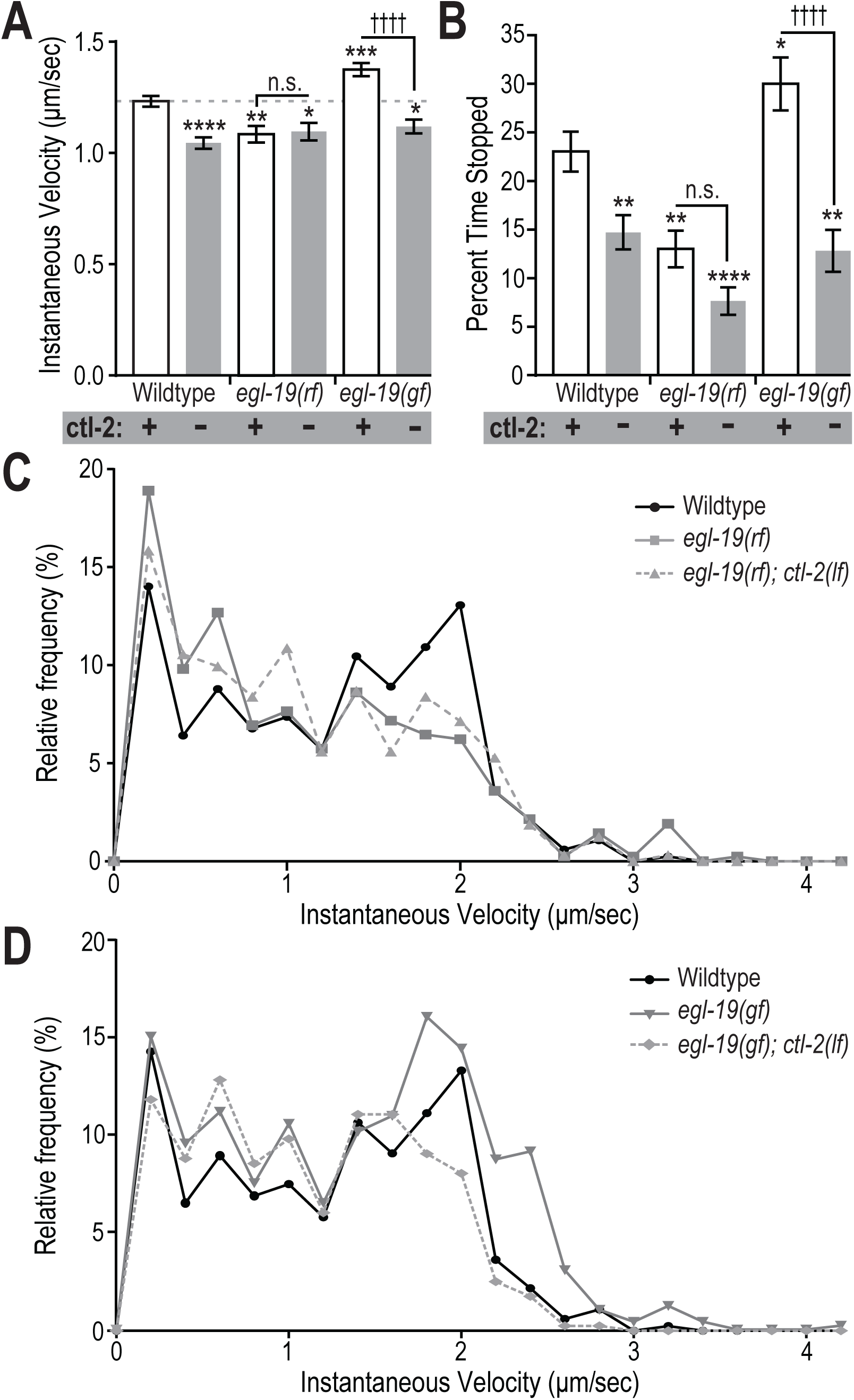
ROS-mediated decrease in GLR-1 transport velocity and stopping is due to effects on L-VGCC signaling. A-D) Stop and velocity analysis of transport events from wild-type, *egl-19(rf)* and *egl-19(gf)* with (+, white bars) and without (-, grey bars) the *ctl-2* gene. A) Instantaneous velocity of anterograde transport events in each group (n≥44 events, n.s.=not significant, *: p<0.05, **: p=0.0035, ***: p=0.0005, ****: p<0.0001 compared to wild-type containing *ctl-2*; ††††: p<0.0001 compared to *egl-19(gf)* alone). B) Percent of time each GLR-1 vesicles spent stopped each group (n≥44 events, *: p=0.044, **: p≤0.009, ****: p<0.0001 compared to wild-type containing *ctl-2*; ††††: p<0.0001 compared to *egl-19*(gf) alone). C and D) Frequency distribution of instantaneous velocity (binned every 0.2 μm/sec) of anterograde transport events for *egl-19(rf)* single and double mutants (C) as well as *egl-19(gf)* single and double mutants (D) in comparison to wild-type containing *ctl-2*.

